# Decoy effect in shoaling decision making in zebrafish

**DOI:** 10.1101/2022.08.31.506011

**Authors:** Abhishek Singh, Kajal Kumari, Sanya Kalra, Shubhi Pal, Bittu Kaveri Rajaraman

## Abstract

The decoy effect, a bias in choice between two options when a third, inferior option is introduced, has been observed across various organisms, from slime molds to humans. In this study, we investigated whether zebrafish (*Danio rerio*), a widely used biological model organism, exhibit the decoy effect in their shoaling group choices, specifically examining this effect by varying only shoal size. Using spatial trajectory analysis of freely swimming zebrafish interacting with conspecifics in adjacent display tanks, we tested how shoaling decisions varied between dichotomous (two-option) and trichotomous (three-option) choice sets. Our experiments compared preferences for 4 versus 2 and 6 versus 3 shoal sizes, with a single fish serving as a decoy in the trichotomous sets. The results revealed sex-specific differences in the decoy effect: males exhibited a shift in preference only in the Trichotomous-first condition, where prior exposure to the decoy led to a significant preference for the larger shoal. In contrast, females displayed the decoy effect exclusively in the Dichotomous-first condition, shifting from a significant preference for the larger shoal to showing no clear preference in the presence of the decoy. Notably, our findings demonstrate that the decoy effect can occur even when studied unidimensionally, with both sex and the order of presentation influencing zebrafish shoaling preferences. These results offer new insights into decision-making processes and highlight the importance of considering order effects in studies of choice behavior, with potential methodological implications for research in other species.

## Introduction

Animals in the wild consistently make decisions that enhance their survival. Living in larger groups offers several advantages, including reduced predation risk through the dilution effect, increased foraging efficiency, improved reproductive success, and better territorial defense (Pitcher 1993; Krause and Ruxton 2002; HR 1984). Larger shoals, while offering protection, also become more conspicuous to predators, increasing the risk of detection. Individuals that differ from the group may face even greater danger due to the ‘oddity effect’ (Landeau and Terborgh 1986; Cattelan and Griggio 2020). Furthermore, increased group size leads to heightened competition, limiting access to resources like food and mates for each member (Grand and Dill 1999; Kent, Holzman, and Genin 2006). This trade-off between the benefits and risks of group living influences shoaling preferences across species. Zebrafish, like many other species, prefer to associate with a shoal rather than remaining isolated (Etinger, Lebron, and Palestis 2009a; Ariyasiri et al. 2019). Among the two sexes, females are known to exhibit a preference for larger shoals (Snekser et al. 2010; Velkey et al. 2022). Shoal size decisions are typically associated with numerical competence and have been observed to adhere to Weber’s law, where the perceived difference between shoal size sets, or numerical sets in general, is represented on a logarithmic scale rather than linearly (Seguin and Gerlai 2017; Gomez-Laplaza and Gerlai 2016; Miletto Petrazzini et al. 2016). A typical shoaling preference test consists of a three-tank setup, where a focal fish in the central tank is presented with two shoal size options in adjacent tanks. The fish’s preference is inferred from the time spent or frequency of visits near the zones adjacent to the display tanks (Lucon-Xiccato et al. 2017; Seguin and Gerlai 2017). Recently, Reding & Cummings (2019) extended the shoaling preference test in mosquitofish to explore multi-alternative shoaling choices and infer rationality in shoal preferences (Reding and Cummings 2019).

The normative principles of economic rationality define individuals as behaving rationally when they make consistent choices that maximize their utility. Such consistent choices can be represented through a stable ordinal utility function that maximizes ‘expected utility’, from which axiomatic principles of transitivity and independence of irrelevant alternatives (IIA) follow (Glimcher and Fehr 2013). Transitivity is complied with if, in a choice set of A, B, and C, an individual who has a stable preference of A over B, and B over C, always prefers A over C. IIA refers to the consistency of preference of A over B regardless of the presence of other items in the choice set (Ray 1973). Specifically, Luce’s choice axiom of constant ratios IIA(L) is upheld when the ratio of the probabilities of choosing A and B in the choice set A, B remains constant even with a larger choice set A, B, C (Luce 1959).

Transitivity and IIA are often reported to be violated by humans (Tversky and Simonson 1993; Trueblood, Brown, et al. 2013; Dumbalska et al. 2020) and several other animals (Latty and Beekman 2011; Morgan et al. 2012; Thomas A Waite 2001; Louie, Khaw, and Glimcher 2013). These violations of rationality largely stem from the context dependence of decision making and suggest that the brain may calculate value relative to context, or through a process of comparison between choice attributes, across available options (Vlaev et al. 2011). Alternatively, relative value information between choice options may be ignored if the animal relies on innate or learned non-compensatory decision-making strategies (Latty and Trueblood 2020). On the other hand, models of value-first decision making, posit an absolute context independent process of valuation underlying rational choices, where options have stable values represented in a cognitive utility scale (Vlaev et al. 2011). IIA refers to one specific aspect of context that can change preferences between two options: the availability of further options-known as distractors or decoys-in the choice set. These context effects are largely reported in studies where the options differ in two or more attribute dimensions with the decoy asymmetrically placed such that its effect dominates relative to one of the attributes, which might result in context specific interactions between the psychophysical value functions of the attributes (Nachev et al. 2017; Morgan et al. 2012; Rivalan, Winter, and Nachev 2017; Bateson, Healy, and Hurly 2003; Bateson, Healy, and Hurly 2002; Parrish, Evans, and Beran 2015; Hu and Yu 2014; Latty and Beekman 2011; Latty and Trueblood 2020; Shafir, Tom A Waite, and Smith 2002; Trueblood and Pettibone 2017). In our knowledge, apart from one study (Morgan et al. 2012), the decoy effect has rarely been reported with options varying along a single dimension of attributes.

Among biological model organisms, the decoy effect (IIA) has only been explored in mice (Rivalan, Winter, and Nachev 2017) and *C. elegans* (Cohen et al. 2019). In C. elegans, chemotaxis assays revealed a decoy effect when choices were processed through an asymmetric neural circuit, while no decoy effect was observed in mice, at least along the single reward dimension of water volume and probability of dispensing drinking water. Zebrafish, a larger and more complex vertebrate model system than *C. elegans*, is similarly equipped with neural imaging and electrophysiological tools, and is also amenable to genetic and pharmacological manipulation, yet remains relatively unexplored. In this study, we investigate the decoy effect in adult zebrafish *(Danio rerio)* using a multi-alternative shoal size choice task. The experimental setup consisted of a central transparent tank surrounded by four display tanks, allowing the focal fish to observe and interact with multiple shoals.We examined this effect unidimensionally, varying only the shoal size of the presented options. The dichotomous comparisons involved shoals of 4 versus 2 fish and 6 versus 3 fish, maintaining the same ratio difference of 0.5 between the options, but differing in absolute shoal size. A third option, a single fish, was included as a decoy in the trichotomous condition. Shoal choice sets were presented in two orders: dichotomous-first and trichotomous-first, for both sexes of the focal fish. This design allowed us to disentangle the effects of the decoy, presentation order, and sex on shoaling preferences, enabling us to determine whether introducing or removing a decoy alters preferences between two shoal sizes.

## Methods

### Subjects

Adult zebrafish (*Danio rerio*) (age <1 year) population was bought from a local pet store in Daryaganj, New Delhi, India. All fish were maintained in the ZebTec Active Blue - Stand Alone system (Tecniplast, PA, USA) and maintained on a 12:12 light: dark (10 a.m. - 10 p.m.) circadian cycle at 7.50-8.50 pH, 28-30°C and 650-700 µS conductivity at Ashoka University, Sonipat, Haryana, India. Fish were fed ad libitum twice a day with powdered Tetra-Tetramin flakes.

We employed 78 adult zebrafish (40 males and 38 females), which were tested for shoaling in both the choice sets of 4 versus 2 and 6 versus 3, each with the decoy option of 1 fish. There was a gap of two weeks between tests. The fish were individually housed in separate tanks within the ZebTec fish system for 2 weeks before the commencement of the shoaling experiments, allowing them to see and smell each other through a shared water circulation system. Female adults from the same population were randomly chosen as display fish for the shoal choice assay.

### Shoal choice apparatus

The shoaling setup comprised a cylindrical acrylic focal tank (24cm diameter, 4mm thickness) within a larger square glass main tank (54cm x 54cm x 31 cm)(Figure 1B). Rectangular display tanks (27cm x 10cm x 30cm) were attached to the main tank, positioned 4cm away from the focal tank. All joints were sealed with DOWSILTM GP silicone sealant. The main tank rested on a raised platform, with its outer faces covered in opaque paper to prevent external visual cues. Laminated black paper covered the perpendicular faces of the display tanks to prevent interaction between display fish.

**Figure 1.**
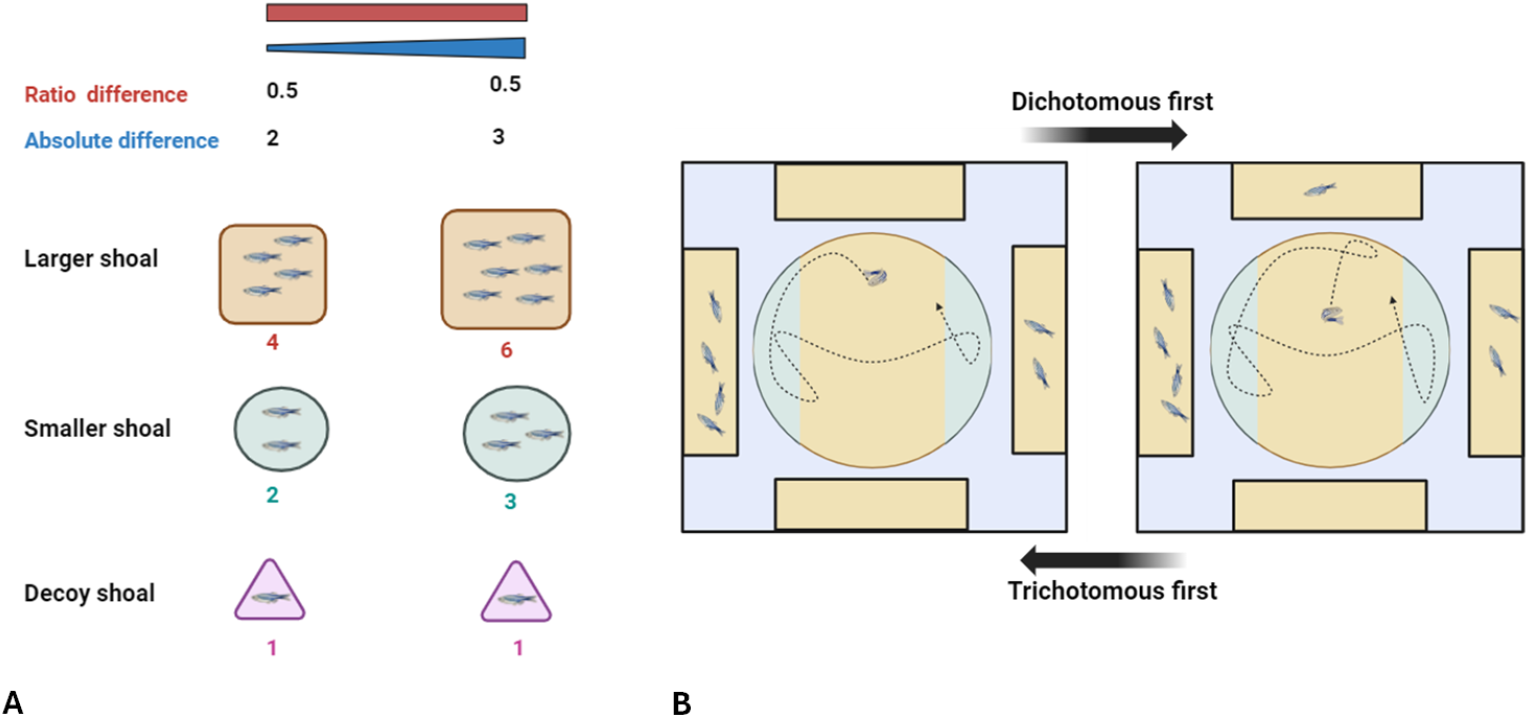
**A**. The shoal choice sets of 4 versus 2 (decoy 1 fish) and 6 versus 3 (decoy 1 fish) differed in absolute shoal size change of 2 and 3, respectively, while maintaining the same ratio difference of 0.5 **B**. Fish movements were recorded for 5 minutes per trial as it navigated the focal tank facing display tanks with shoal choice sets presented in either Dichotomous-first or Trichotomous-first order. Time spent in sectors facing the display tanks was calculated using trajectory data.

An LED light strip (Mufasa Copper Non-Waterproof LED Strip 5050) was attached to each display tank to provide internal lighting and prevent tripod shadows. A dummy tripod, rising to the same height, approached the existing camera tripod from the opposite end. The entire setup was covered by a 10ft X 10ft black photography cloth curtain to block external visual cues, ensuring the sole light source was from the display tanks. Water filled the space between display and focal tanks to minimize refraction distortion. The background face of the display tanks, behind the display fish, was covered with a green sheet of laminated paper, as per Lucon-Xiccato et al. (2017). We used a GoPro Hero 8 camera (1080p|30fps|linear) for these recordings.

### Experimental design

We analyzed the trajectories of adult zebrafish within a central circular tank as they shoaled alongside groups of zebrafish presented as choice options in display tanks. The aim was to infer their relative preference for larger shoals (4 or 6 fish) over smaller shoals (2 or 3 fish), both with and without the decoy option of a single fish (Figure 1). Before each trial, a cylindrical opaque plastic sheet was placed between the focal and display tanks to visually isolate the focal fish from the display fish.

To control for potential biases between different sections of the tank, a no-fish trial was conducted prior to the experimental trials. In this trial, the subject fish was introduced to the focal tank and exposed to empty display tanks after a 2-minute visual isolation. Following the no-fish trial, the subject fish was randomly assigned to either the Dichotomous-first or Trichotomous-first condition. In both conditions, the display fish were introduced into the display tanks while the focal fish remained visually isolated. Each trial, whether dichotomous or trichotomous, lasted for 5 minutes, during which video recordings of the focal fish’s shoaling responses were captured. This was followed by a 2-minute resting period, during which the display fish were replaced for the next trial.

To randomize the experimental conditions, the *Sample()* function in R was used to first randomly deter-mine the axis of presentation of the dichotomous choice (relative to the camera), followed by randomizing the side of the larger shoal presentation, and finally the side of the third choice (for trichotomous trials). The order of testing for each choice set was randomized, and each fish was randomly assigned to either the 4 versus 2 (decoy 1) or 6 versus 3 (decoy 1) choice set, with a 2-week gap between tests.

In the 4 versus 2 (decoy 1) choice set, 19 males and 19 females experienced the Trichotomous-first condition, while in the 6 versus 3 (decoy 1) choice set, 20 males and 20 females were tested in the same condition, out of a total sample of 40 males and 38 females.

### DeepLabCut based fish tracking

All videos were cropped to a fixed duration of 5 minutes, starting from the moment the sheet covering the focal tank was raised, using the Python-based video editing tool Moviepy. The videos were then processed using DeepLabCut (DLC), a Python-based pose estimation package (Mathis et al. 2018), to track the head, body, and tail of the fish. A subset of videos was randomly selected to generate training frames, manually marking pre-defined body parts. The default Artificial Neural Network ResNet 50 was trained for more than 400,000 iterations. Labeled videos were generated to verify tracking efficiency. Tracking data for each body part were converted and stored in .csv format. We chose the body part which was most efficiently tracked by inspecting the labeled tracked videos, which in most cases was the ‘body’ point, rather than the head or tail.

### Analysis

The trajectory data obtained from the DLC tracking was processed using a custom R script DLC-Analyzer (Sturman et al. 2020) for the time spent in each sector data. The preference index for the larger shoal was calculated using the time spent in each sector data as :

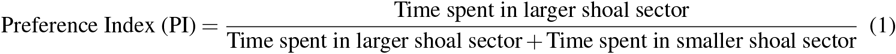

We analyzed the preference index data in two steps: first, using a generalized linear mixed model (GLMM) for direct comparisons, and second, a one-sample t-test to compare the results against chance.

For the direct comparisons, we used a GLMM with the Preference Index for the larger shoal as the response variable. The explanatory variables included Shoal Presentation (‘Shoals’, Dichotomous or Trichotomous), Order of Presentation (‘Order’, Dichotomous First or Trichotomous First), and Sex(‘Sex’, Males or Females), with Individual ID included as a random factor. The analysis was conducted separately for the 4 versus 2 (decoy 1) and 6 versus 3 (decoy 1) choice sets. Interactions between all explanatory variables were included to capture the effects of the decoy option. Since the preference index ranged between 0 and 1, we applied a GLMM with a beta family distribution and a logit link function using the ‘glmmTMB’ package in R (Brooks et al. 2017). We checked for dispersion of the residuals using the ‘DHARMa’ package in R and found no significant issues (Hartig 2018) (see Supplementary Figures 1 and 2). For pairwise comparisons, we performed a post-hoc test with Tukey correction using the contrast package in R.

In the second step, we conducted one-sample t-tests to determine whether preferences deviated from chance (preference index of 0.5). These tests were used to assess the preferences for each combination of Shoal Presentation, Order, and Sex. The data were normally distributed, and since each test addressed independent and parallel hypotheses without influencing one another, the alpha level for all tests was maintained at 0.05.

To visually inspect trajectory densities alongside preference index data based on time spent in zones (1), we used 2D Kernel Density Estimates (KDE) with a Gaussian kernel. Since shoals were presented in a randomized manner across the four display tanks surrounding the central cylindrical tank, all trajectories were first rotated using the ‘SMoLR’ (Paul et al. 2019) package in R to align along the 4-2/6-3 axis (top-down orientation). These aligned trajectories were then combined to create a single, overlapping population density plot using the ‘spatstat’ (Baddeley, Rubak, and Turner 2015) package in R.

### Ethical Note

All procedures involving animals were conducted in accordance with the guidelines established by the Committee for Control and Supervision of Experiments on Animals (CCSEA), adhering to both national and institutional ethical standards for the care and use of animals. Ethical approval was obtained from Ashoka University’s Institutional Animal Ethics Committee (approval no. ASHOKA/IAEC/2/2022/6).

## Results

No zone bias was observed in the time spent by male and female populations in each sector during the initial no-fish trials (see Supplementary Figures 3 and 4).

No significant differences were found in direct comparisons between all combinations of Shoal Presentation, Sex, and Order of Presentation (see Supplementary Table 5 and 6, Tukey post-hoc test; all p-values > 0.05) for both the 4 versus 2 (decoy 1) and 6 versus 3 (decoy 1) choice sets. The presentation of the trichotomous shoal (with the decoy) significantly decreases the preference for the larger shoal (4 fish) at the population level in the 4 versus 2 (decoy 1) choice set (see Supplementary Table 3, Wald Chi-squared = 5.21, p < 0.05), averaging over levels of sex and order.

To test for shifts in relative preference for the larger shoal against chance, we further investigated these effects using one-sample t-tests against a chance-level Preference Index of 0.5 for each combination of Sex, Presentation Order, and Shoaling Condition. Additionally, we employed 2D Kernel Density Estimation (KDE) plots to visualize and detect subtle spatial shifts in preference.

In the 4 versus 2 (decoy 1) choice set, as illustrated in Figure 2, males did not exhibit a baseline preference for the larger shoal (4) over the smaller shoal (2) in the Dichotomous-first condition. The introduction of a decoy in the Trichotomous choice set did not significantly alter preferences; however, the KDE plot suggested a shift toward the smaller shoal (2). In the Trichotomous-first condition, there was no initial preference for the larger shoal in the Trichotomous choice set, but a significant preference for the larger shoal emerged in the subsequent Dichotomous choice set (Figure 2: Mean = 0.65, SEM = 0.06, t(18) = 2.27, p < 0.05), as reflected in the trajectory density plot. Females initially preferred the larger shoal in the Dichotomous-first condition (Figure 2: Mean = 0.70, SEM = 0.06, t(18) = 2.92, p < 0.01), but this preference disappeared in the following Trichotomous choice set. In the Trichotomous-first condition, no significant preference was observed in either choice set.

**Figure 2.**
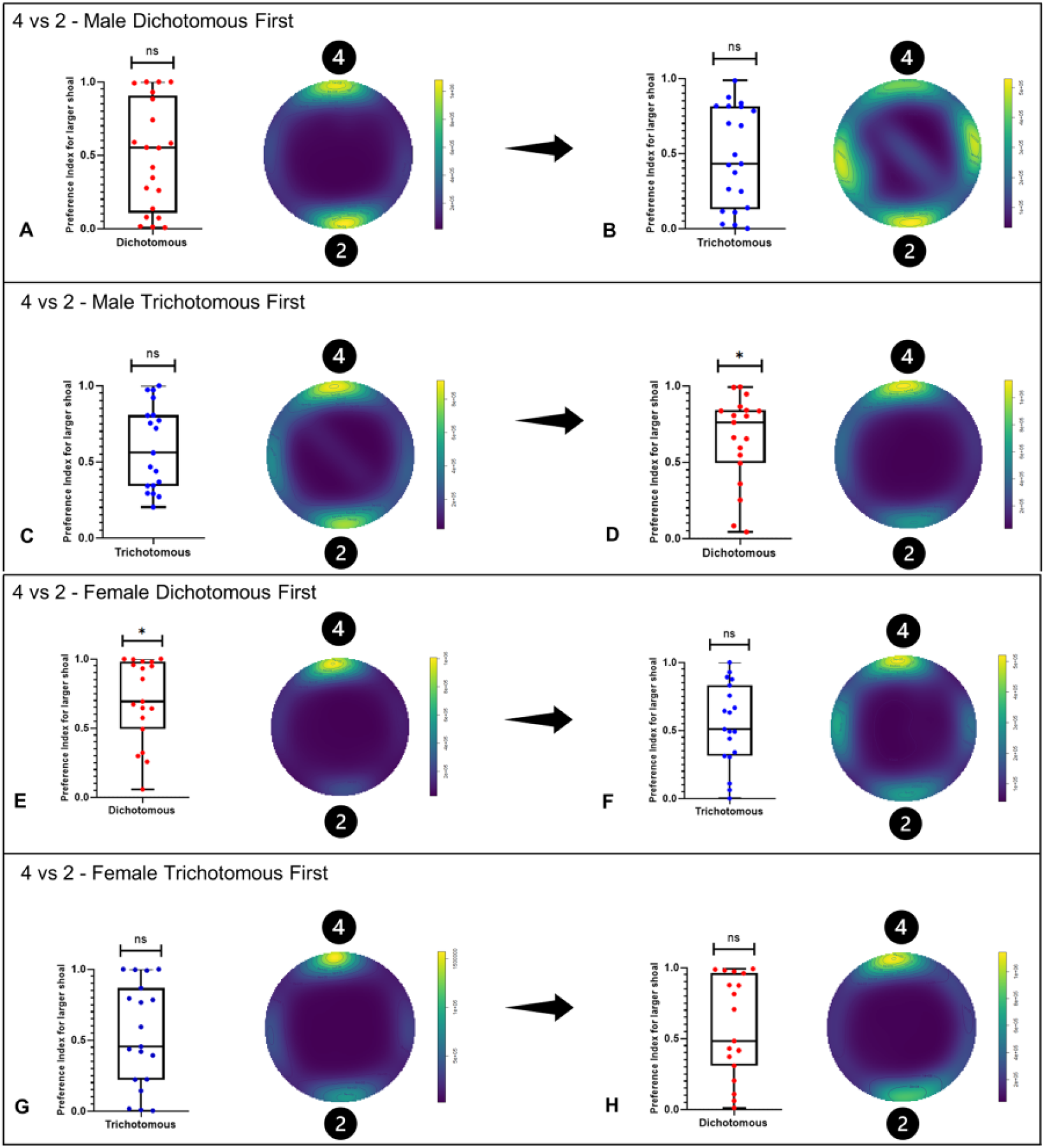
Preference indices for the larger shoal (4 versus 2) for both males and females, shown for Dichotomous and Trichotomous choice sets in both Dichotomous-first and Trichotomous-first conditions, with values ranging from 0 to 1 for comparison against the chance level (0.5). Each panel (A-H) also includes 2D Gaussian KDE trajectory plots for the respective choice sets.

In the 6 versus 3 (decoy 1) choice set, males did not show a preference for the larger shoal (6) over the smaller shoal (3) in the Dichotomous-first condition. The introduction of a decoy led to a shift towards the smaller shoal (3), although this change was not statistically significant when comparing relative preference against chance. A significant relative preference for the larger shoal (6) emerged in the Dichotomous choice set of the Trichotomous-first condition (Figure 3: Mean = 0.63, SEM = 0.05, t(19) = 2.49, p < 0.05). The shift in the direction of the decoy effect with changes in the presentation order was evident in the trajectory density plots. Females showed only a weak and nonsignificant baseline preference for the larger shoal (Figure 3: Mean = 0.62, SEM = 0.07, t(19) = 1.70, p = 0.10) in the Dichotomous-first condition and did not exhibit any significant preference in either choice set of the Trichotomous-first condition. For all statistical values from the one-sample t-test comparisons, see Table 1.

**Figure 3.**
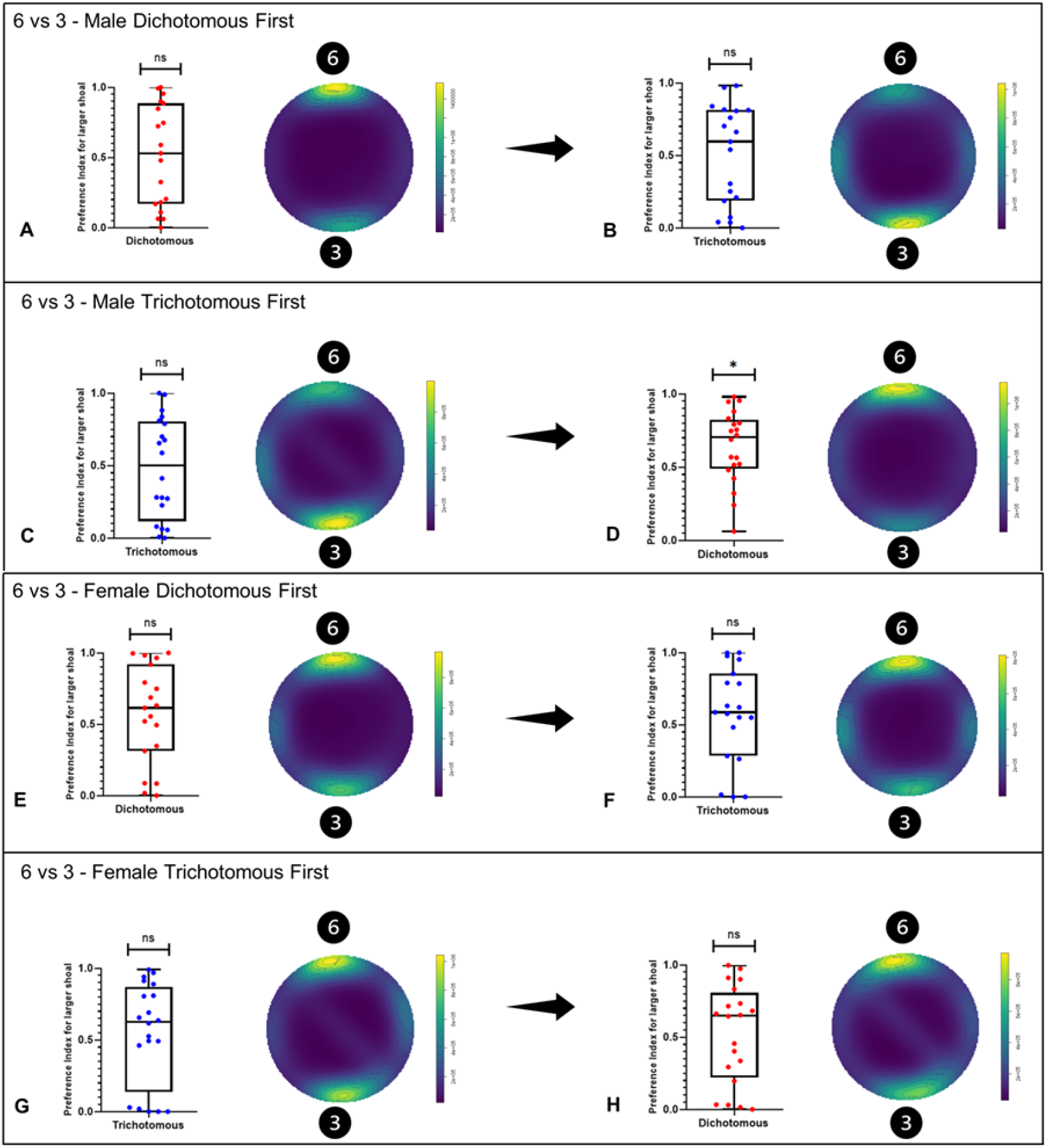
Preference indices for the larger shoal (6 versus 3) for both males and females, shown for Dichotomous and Trichotomous choice sets in both Dichotomous-first and Trichotomous-first conditions, with values ranging from 0 to 1 for comparison against the chance level (0.5). Each panel (A-H) also includes 2D Gaussian KDE trajectory plots for the respective choice sets.

**Table 1.** Results from One sample t-test comparisons for each condition.

| 4 versus 2 (decoy 1) |  |  |  |  | 6 versus 3 (decoy 1) |  |  |  |  |
| --- | --- | --- | --- | --- | --- | --- | --- | --- | --- |
|  | Mean | +/- SEM | t | p |  | Mean | +/- SEM | t | p |
| **Males dichotomous first (N=20)** |  |  |  |  | **Males dichotomous first (N=19)** |  |  |  |  |
| Dichotomous | 0.49 | 0.08 | -0.03 | 0.97 | Dichotomous | 0.51 | 0.08 | 0.16 | 0.86 |
| Trichotomous | 0.47 | 0.07 | -0.35 | 0.72 | Trichotomous | 0.50 | 0.07 | 0.06 | 0.95 |
| **Males trichotomous first (N=19)** |  |  |  |  | **Males trichotomous first (N=20)** |  |  |  |  |
| Dichotomous | 0.65** | 0.06 | 2.27 | 0.03 | Dichotomous | 0.63** | 0.05 | 2.49 | 0.02 |
| Trichotomous | 0.59 | 0.06 | 1.49 | 0.15 | Trichotomous | 0.48 | 0.07 | -0.25 | 0.80 |
| **Females dichotomous first (N=19)** |  |  |  |  | **Females dichotomous first (N=19)** |  |  |  |  |
| Dichotomous | 0.70*** | 0.06 | 2.92 | 0.009 | Dichotomous | 0.62* | 0.07 | 1.70 | 0.10 |
| Trichotomous | 0.54 | 0.06 | 0.60 | 0.55 | Trichotomous | 0.58 | 0.07 | 1.11 | 0.28 |
| **Females trichotomous first (N=19)** |  |  |  |  | **Females trichotomous first (N=20)** |  |  |  |  |
| Dichotomous | 0.57 | 0.08 | 0.96 | 0.34 | Dichotomous | 0.52 | 0.07 | 0.31 | 0.75 |
| Trichotomous | 0.53 | 0.08 | 0.38 | 0.70 | Trichotomous | 0.54 | 0.07 | 0.59 | 0.56 |

\*p<0.1 \*\*p<0.05 \*\*\*p<0.01

## Discussion

Our study provides the first evidence of the decoy effect in shoal size choice among zebrafish (*Danio rerio*), a neurobiologically and genetically tractable model organism. We demonstrate that both sex and the order of presentation influence this decoy effect. To investigate these effects, we employed a combination of methods: generalized linear mixed models for direct comparisons, one-sample t-tests to assess shifts against chance, and 2D Kernel Density Estimation (KDE) plots to visualize subtle shifts in trajectory density.

In the direct comparison using GLMM, the introduction of a decoy had a significant effect on shoal preference only in the 4 versus 2 (decoy 1) choice set at the population level. However, when tested against chance (0.5 preference index) and visualized through trajectory density plots (2D KDE), we observed notable shifts in preference based on sex and presentation order in both 4 versus 2 and 6 versus 3. While the traditional framework to test for decoy effect focuses on testing violations of the Independence of Irrelevant Alternatives (IIA) through direct comparisons, this approach may miss small, yet ecologically important effects.

A preference for the larger shoal was observed only among females in the 4 versus 2 choice set. In females, there was a shift from significantly preferring the larger shoal (4) in the Dichotomous choice set to not preferring any shoal in the Trichotomous choice set of the Dichotomous-first condition. Females showed only a weak (non-significant) preference for the shoal size of 6 over 3. In both the 4 versus 2 (decoy 1) and 6 versus 3 (decoy 1) choice sets, no effect was observed in the Trichotomous-first condition of the preference index data. The 2D KDE plots confirmed that decoy presentation had no significant effect in the Trichotomous-first condition for either choice set. However, in the Dichotomous-first condition, the introduction of the decoy led to a noticeable shift in trajectory density, reducing the strong preference for the larger shoal, consistent with the time spent data.

Interestingly, males who initially showed no preference for the larger shoal in the Dichotomous-first condition developed a significant preference for the larger shoal in both the 4 versus 2 and 6 versus 3 choice sets after being exposed to the decoy option in prior trials under the Trichotomous-first condition. The 2D KDE trajectory plots confirmed a notable shift in shoal preference depending on the order of decoy presentation. When the decoy was presented first, followed by the Dichotomous choice set, there was a shift toward the larger shoal. Conversely, in the Dichotomous-first condition, a shift toward the smaller shoal was observed, although this shift was not statistically significant (against chance) in the time spent data and was only noticeable in the trajectory densities from the 2D KDE plots.

The shift observed in males, where a preference for the larger shoal emerged after prior exposure to the decoy option, resembles the design of phantom decoy tasks, commonly used in the field (Orlando et al. 2023; Marini et al. 2023), where a third option is presented but unavailable at the time of choice. The change in preference among females—from significantly selecting the larger shoal (against chance) in the Dichotomous-first trial to not choosing it in the subsequent Trichotomous trial—could be attributed to the “random dilution effect” (Bateson, Healy, and Hurly 2002), where a proportion of choices are randomly allocated between options. A third option may absorb these even with no change of bias between the two main options, thereby diluting the masking effect of the random noise on the existing choice bias and appearing to change the relative preference (Bateson 2002).

Our findings of sex-based differences in shoal size preference align with previous reports of similar patterns in shoal discrimination (Ruhl and McRobert 2005; Etinger, Lebron, and Palestis 2009b; Way et al. 2015; Velkey et al. 2022). However, many other studies on shoal discrimination have not specifically tested for sex-based differences in shoaling behavior (Seguin and Gerlai 2017; Sheardown et al. 2022). Additionally, most shoal choice studies employ a different setup—a rectangular three-tank apparatus. Unlike these previous studies, our results reveal clear sex-based differences in preference for larger shoals and notable shifts in behavior when a third, inferior (decoy) option is introduced. Moreover, the order of decoy presentation had a differential effect on males and females. These differences could be driven by distinct evolutionary pressures influencing shoal choice between the sexes (Snekser et al. 2010; Magurran and Garcia 2000). In the 6 versus 3 shoal choice task, males showed no strong preference, while females exhibited a weak (non-significant) preference for the larger shoal. In contrast, females displayed a significant preference for the larger shoal in the 4 versus 2 task. This contrasts with findings by Seguin & Gerlai (2017) (Seguin and Gerlai 2017), who reported a significant population-level preference for the larger shoal in the 6 versus 3 task. The discrepancy may be attributed to differences in experimental conditions, such as the strains of zebrafish used or the introduction of a perforated divider in their study, which allowed both olfactory and visual cues that could have influenced shoaling decisions.

The influence of order effects in decoy studies is often overlooked or underreported. One recent study (Orlando et al. 2023) allowed swamp wallabies to freely choose between dichotomous and trichotomous options in the wild, revealing a significant order effect. However, the order was not the primary variable of interest, and the authors discussed the main effects without addressing the specific role of order in detail. Phantom decoy tasks introduce a specific order by presenting the decoy first and then measuring shifts in preference without it, resembling the Trichotomous-first condition in our study. However, this approach only allows for the examination of preferences in one specific order (Trichotomous-first), limiting the ability to explore how preferences might change based solely on the order of presentation of the decoy option. Our study presents the first evidence, to our knowledge, that the direction of the decoy effect is influenced by the order of presentation and is also sex-specific. Our results underscore that the decoy effect should not be studied exclusively using a single order of presentation, as illustrated by the phantom-decoy paradigm. Evaluating the decoy effect without accounting for the order of presentation may lead to misinterpretations of its isolated effects.

Context effects in decision-making have been reported in the fish literature, largely regarding multi-attribute mate choice by females, for example, in the peacock blenny (Salaria pavo) (Locatello, Poli, and Rasotto 2015) and the green swordtail (Xiphophorus helleri) (Royle, Lindström, and Metcalfe 2008). The investigation of independence of irrelevant alternatives (IIA) in shoal size choice decisions has only been conducted in mosquitofish (Gambusia affinis), where no violation of the constant ratio rule was found (Reding and Cummings 2019). However, like most studies, they performed a direct comparison of relative preference using a linear model to rule out the decoy effect. We adopted a similar experimental design in our shoal size choice task to avoid the replication crisis seen in human decoy effect studies (Rivalan, Winter, and Nachev 2017; Gluth, Spektor, and Rieskamp 2018; Chau, Kolling, et al. 2014; Chau, Law, et al. 2020), where small variations in the task and stimulus modality may have contributed to the differing decoy effects observed.

Unidimensional violations of IIA have been rarely reported, with only one known instance (Morgan et al. 2012). Such research is crucial due to the challenge of interpreting multi-attribute context effects, where multiple plausible underlying models can lead to the same outcome. Morgan et al. (2012) also demonstrated different decoy effects occurring during single-attribute manipulations for different attributes (Morgan et al. 2012). It is reasonable to expect that different mechanisms mediate context effects in various single-attribute choices, and these need exploration for a comprehensive understanding of decision-making regarding various attributes considered together.

Our study establishes the presence of decoy effects in zebrafish shoaling decisions, with distinct influences of order and sex, all in relation to a single attribute: shoal size. The extensive array of pharmacological, genetic, neurobiological, and behavioral tools available in this system will facilitate future research aimed at comprehensively understanding context effects in this model organism. Moreover, these tools will enable the examination of the underlying mechanisms behind sex-specific rationality in contextual decision-making. Furthermore, these results significantly contribute to the existing decoy effect literature by focusing on decoy effects at single dimensions and highlighting their absence in investigations and modeling of multi-alternative, multi-attribute decoy effects.

## Acknowledgements

This research was supported by the Ashoka University Annual Faculty Research grant provided to BKR and the infrastructure provided by the Research Office at Ashoka University. AS would like to thank his Ph.D. Research Committee members Aurnab Ghose, Krishna Melnattur and Joby Joseph for feedback on the experiments and analysis. The authors would like to thank Shivani Krishna, Sudipta Tung and LS Shashidhara for feedback on the manuscript.

## Author contributions statement

The study was designed by AS, KK and BKR; the apparatus was made by AS and KK; data collection was carried out by AS, SK, and KK; Video and trajectory analysis was performed by AS,SK and SP; data analysis was carried out by AS; and the manuscript was written by AS and BKR.

## Data Availability

Supplementary information and raw data files that support the findings of this study are openly available on Figshare at https://doi.org/10.6084/m9.figshare.20745865.

