## Supplementary Information for "Decoy effect in shoaling decision making in zebrafish"

Model : Pref_index ~ Shoals*Sex*Order+ (1|ID) , family = beta_family(link = "logit")

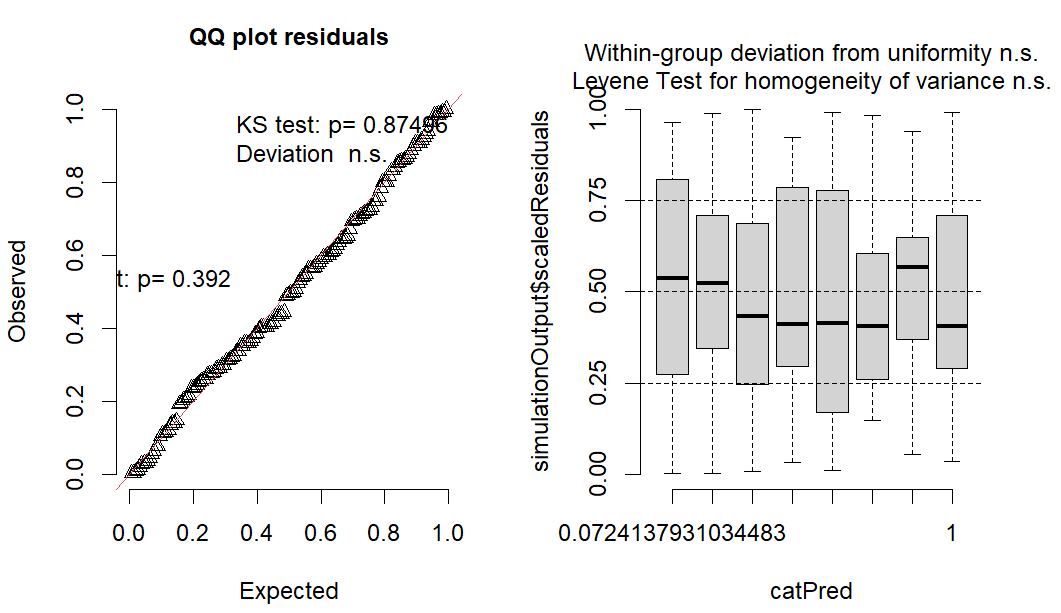

**Figure 1** – Q-Q and Residual vs Predicted plots for 4 versus 2 decoy 1 beta (logit link) GLMM

Model : Pref_index ~ Shoals*Sex*Order+ (1|ID) , family = beta_family(link = "logit")

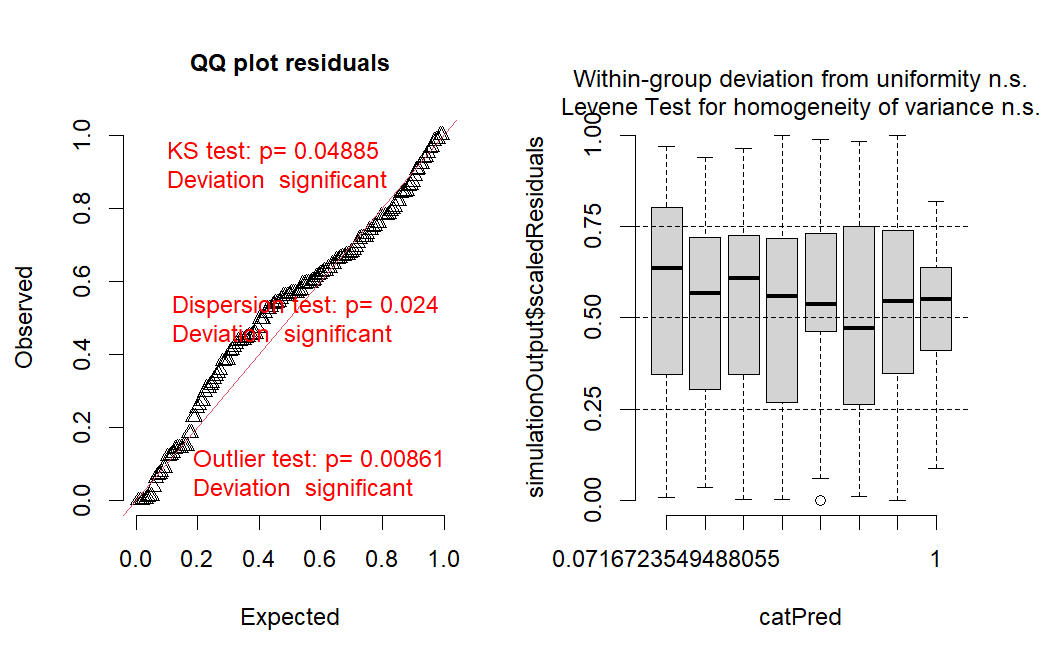

model.

**Figure 2** – Q-Q and Residual vs Predicted plots for 6 versus 3 decoy 1 beta (logit link) GLMM model.

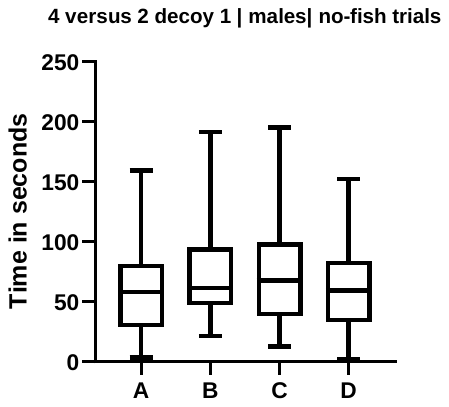

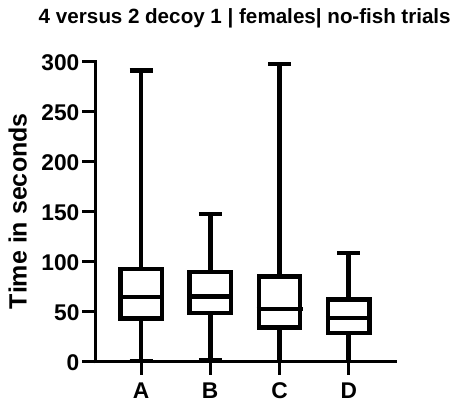

**Figure 3** - Time spent in each sector during the no-fish trials of the 4 versus 2 (decoy 1) choice set, demonstrating no significant zone bias.

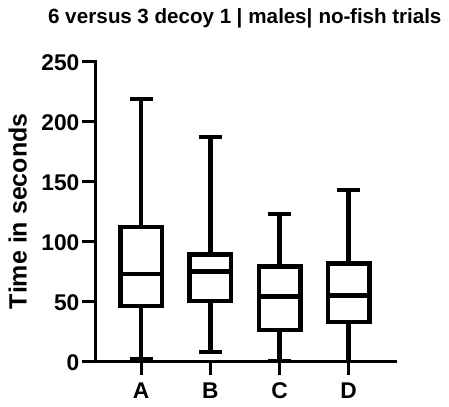

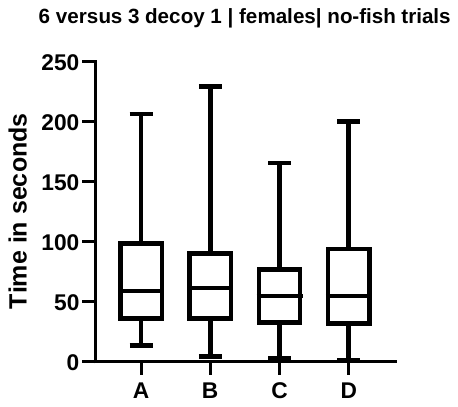

**Figure 4** - Time spent in each sector during the no-fish trials of the 6 versus 3 (decoy 1) choice set, demonstrating no significant zone bias.

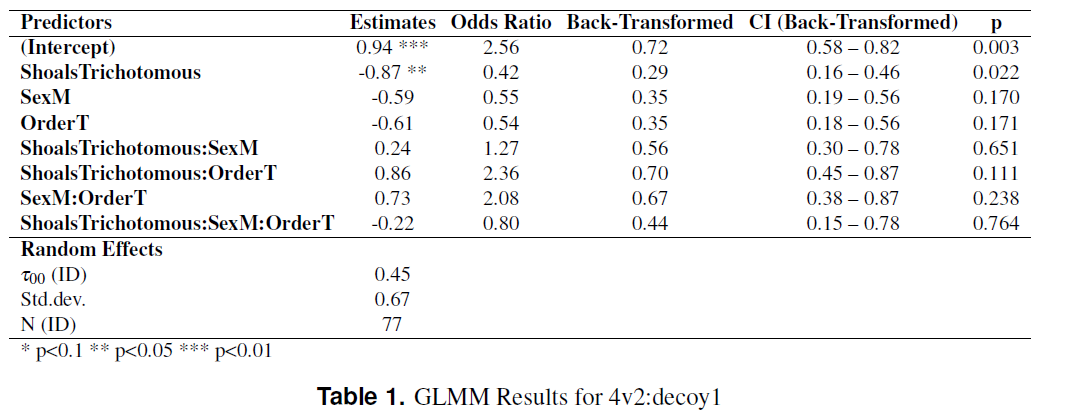

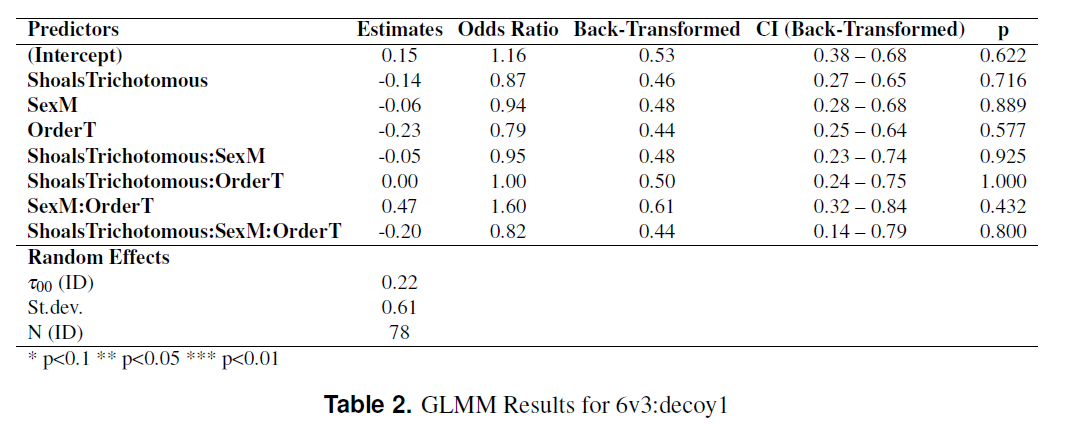

Model : Pref_index ~ Shoals*Sex*Order+ (1|ID) , family = beta_family(link = "logit")

| Response : Preference Index |  |  |  |
| --- | --- | --- | --- |
|  | Chisq | df | p |
| (Intercept) | 8.86 | 1 | **0.002** |
| ShoalsTrichotomous | 5.21 | 1 | **0.022** |
| SexM | 1.88 | 1 | 0.17 |
| OrderT | 1.87 | 1 | 0.17 |
| ShoalsTricho:SexM | 0.20 | 1 | 0.65 |
| ShoalsTricho:OrderT | 2.54 | 1 | 0.11 |
| SexM:OrderT | 1.39 | 1 | 0.23 |
| ShoalsTricho:SexM:OrderT | 0.09 | 1 | 0.76 |

**Table 3 :** Analysis of Deviance Table (Type III Wald chisquare tests) (4 vs 2 decoy 1 )

Model : Pref_index ~ Shoals*Sex*Order+ (1|ID) , family = beta_family(link = "logit")

| Response : Preference Index |  |  |  |
| --- | --- | --- | --- |
|  | Chisq | df | p |
| (Intercept) | 0.24 | 1 | 0.622 |
| ShoalsTrichotomous | 0.13 | 1 | 0.715 |
| SexM | 0.01 | 1 | 0.889 |
| OrderT | 0.31 | 1 | 0.577 |
| ShoalsTricho:SexM | 0.08 | 1 | 0.924 |
| ShoalsTricho:OrderT | 0.00 | 1 | 1.000 |
| SexM:OrderT | 0.61 | 1 | 0.432 |
| ShoalsTricho:SexM:OrderT | 0.06 | 1 | 0.799 |

**Table 4 :** Analysis of Deviance Table (Type III Wald chisquare tests) (6 vs 3 decoy 1 )

**Table 5 :** Results of pairwise comparisons from Post Hoc Tukey test for 4 versus 2 (decoy 1) choice set

Model : Pref_index ~ Shoals*Sex*Order+ (1|ID) , family = beta_family(link = "logit")

| contrast | estimate | SE | df | z.ratio | p.value |
| --- | --- | --- | --- | --- | --- |
| Dichotomous F D - Trichotomous F D | 0.87856 | 0.385 | Inf | 2.285 | 0.3019 |
| Dichotomous F D - Dichotomous M D | 0.5955 | 0.434 | Inf | 1.372 | 0.8701 |
| Dichotomous F D - Trichotomous M D | 1.232 | 0.444 | Inf | 2.772 | 0.1021 |
| Dichotomous F D - Dichotomous F T | 0.61187 | 0.447 | Inf | 1.369 | 0.8716 |
| Dichotomous F D - Trichotomous F T | 0.62937 | 0.441 | Inf | 1.428 | 0.8444 |
| Dichotomous F D - Dichotomous M T | 0.46829 | 0.449 | Inf | 1.044 | 0.9677 |
| Dichotomous F D - Trichotomous M T | 0.47347 | 0.448 | Inf | 1.056 | 0.9655 |
| Trichotomous F D - Dichotomous M D | -0.28306 | 0.428 | Inf | -0.661 | 0.9979 |
| Trichotomous F D - Trichotomous M D | 0.35344 | 0.438 | Inf | 0.807 | 0.9928 |
| Trichotomous F D - Dichotomous F T | -0.2667 | 0.443 | Inf | -0.602 | 0.9989 |
| Trichotomous F D - Trichotomous F T | -0.24919 | 0.434 | Inf | -0.574 | 0.9992 |
| Trichotomous F D - Dichotomous M T | -0.41027 | 0.446 | Inf | -0.92 | 0.9843 |
| Trichotomous F D - Trichotomous M T | -0.40509 | 0.445 | Inf | -0.911 | 0.9852 |
| Dichotomous M D - Trichotomous M D | 0.6365 | 0.372 | Inf | 1.711 | 0.6804 |
| Dichotomous M D - Dichotomous F T | 0.01637 | 0.434 | Inf | 0.038 | 1 |
| Dichotomous M D - Trichotomous F T | 0.03388 | 0.424 | Inf | 0.08 | 1 |
| Dichotomous M D - Dichotomous M T | -0.12721 | 0.437 | Inf | -0.291 | 1 |
| Dichotomous M D - Trichotomous M T | -0.12203 | 0.435 | Inf | -0.28 | 1 |
| Trichotomous M D - Dichotomous F T | -0.62014 | 0.443 | Inf | -1.398 | 0.8584 |
| Trichotomous M D - Trichotomous F T | -0.60263 | 0.435 | Inf | -1.387 | 0.8636 |
| Trichotomous M D - Dichotomous M T | -0.76371 | 0.446 | Inf | -1.711 | 0.6799 |
| Trichotomous M D - Trichotomous M T | -0.75853 | 0.445 | Inf | -1.704 | 0.6849 |
| Dichotomous F T - Trichotomous F T | 0.01751 | 0.382 | Inf | 0.046 | 1 |
| Dichotomous F T - Dichotomous M T | -0.14358 | 0.45 | Inf | -0.319 | 1 |
| Dichotomous F T - Trichotomous M T | -0.1384 | 0.449 | Inf | -0.308 | 1 |
| Trichotomous F T - Dichotomous M T | -0.16109 | 0.443 | Inf | -0.364 | 1 |
| Trichotomous F T - Trichotomous M T | -0.15591 | 0.441 | Inf | -0.353 | 1 |
| Dichotomous M T - Trichotomous M T | 0.00518 | 0.395 | Inf | 0.013 | 1 |

**Table 6 :** Results of pairwise comparisons from Post Hoc Tukey test for 6 versus 3 (decoy 1) choice set

Model : Pref_index ~ Shoals*Sex*Order+ (1|ID) , family = beta_family(link = "logit")

| contrast | estimate | SE | df | z.ratio | p.value |
| --- | --- | --- | --- | --- | --- |
| Dichotomous F D - Trichotomous F D | 0.1497 | 0.411 | Inf | 0.364 | 1 |
| Dichotomous F D - Dichotomous M D | 0.0606 | 0.436 | Inf | 0.139 | 1 |
| Dichotomous F D - Trichotomous M D | 0.265 | 0.439 | Inf | 0.604 | 0.9988 |
| Dichotomous F D - Dichotomous F T | 0.239 | 0.429 | Inf | 0.558 | 0.9993 |
| Dichotomous F D - Trichotomous F T | 0.3887 | 0.427 | Inf | 0.911 | 0.9851 |
| Dichotomous F D - Dichotomous M T | -0.18 | 0.439 | Inf | -0.41 | 0.9999 |
| Dichotomous F D - Trichotomous M T | 0.2287 | 0.429 | Inf | 0.533 | 0.9995 |
| Trichotomous F D - Dichotomous M D | -0.0891 | 0.434 | Inf | -0.205 | 1 |
| Trichotomous F D - Trichotomous M D | 0.1153 | 0.438 | Inf | 0.264 | 1 |
| Trichotomous F D - Dichotomous F T | 0.0893 | 0.427 | Inf | 0.209 | 1 |
| Trichotomous F D - Trichotomous F T | 0.239 | 0.425 | Inf | 0.562 | 0.9993 |
| Trichotomous F D - Dichotomous M T | -0.3297 | 0.437 | Inf | -0.754 | 0.9952 |
| Trichotomous F D - Trichotomous M T | 0.079 | 0.428 | Inf | 0.185 | 1 |
| Dichotomous M D - Trichotomous M D | 0.2044 | 0.408 | Inf | 0.501 | 0.9997 |
| Dichotomous M D - Dichotomous F T | 0.1784 | 0.426 | Inf | 0.419 | 0.9999 |
| Dichotomous M D - Trichotomous F T | 0.3281 | 0.423 | Inf | 0.775 | 0.9944 |
| Dichotomous M D - Dichotomous M T | -0.2406 | 0.435 | Inf | -0.553 | 0.9993 |
| Dichotomous M D - Trichotomous M T | 0.1681 | 0.427 | Inf | 0.394 | 0.9999 |
| Trichotomous M D - Dichotomous F T | -0.026 | 0.429 | Inf | -0.061 | 1 |
| Trichotomous M D - Trichotomous F T | 0.1237 | 0.427 | Inf | 0.29 | 1 |
| Trichotomous M D - Dichotomous M T | -0.445 | 0.439 | Inf | -1.013 | 0.9726 |
| Trichotomous M D - Trichotomous M T | -0.0363 | 0.43 | Inf | -0.084 | 1 |
| Dichotomous F T - Trichotomous F T | 0.1497 | 0.388 | Inf | 0.386 | 0.9999 |
| Dichotomous F T - Dichotomous M T | -0.419 | 0.428 | Inf | -0.978 | 0.9776 |
| Dichotomous F T - Trichotomous M T | -0.0103 | 0.419 | Inf | -0.025 | 1 |
| Trichotomous F T - Dichotomous M T | -0.5687 | 0.426 | Inf | -1.333 | 0.8862 |
| Trichotomous F T - Trichotomous M T | -0.16 | 0.417 | Inf | -0.383 | 0.9999 |
| Dichotomous M T - Trichotomous M T | 0.4087 | 0.404 | Inf | 1.011 | 0.9729 |
